# Pontocerebellar Hypoplasia linked mutations of the deadenylase Target of EGR1 (TOE1) impair thermal stability, ribonuclease activity, and oligomerization

**DOI:** 10.1101/2025.09.18.672001

**Authors:** Lizette Zavala, Magdalena Sobien, Cassandra K. Hayne

## Abstract

The Target of EGR1 (TOE1) gene encodes the TOE1 deadenylase, which is essential for the maturation of Pol-II transcribed snRNAs in humans. Over a dozen missense mutations in the TOE1 gene have been linked to Pontocerebellar Hypoplasia Type 7 (PCH7), a rare but serious neurodevelopmental and neurodegenerative disease that leads to early mortality. The biochemical mechanisms for why these PCH7-linked mutations alter TOE1’s biochemical characteristics remains vague. Here, we utilized AlphaFold predicted structures of TOE1 and biochemical characterizations to investigate the impact of selected TOE1 variants on TOE1’s biochemical properties.

We performed characterization of the thermal stability and activity of eleven PCH-linked TOE1 variants and found that eight variants have significantly reduced protein thermal stability and that all but two variants impair TOE1’s ribonuclease activity, particularly its exonuclease activity. Additionally, we found that the F148Y mutation impacts TOE1’s oligomeric state *in vitro* and *in vivo*. Together, these results demonstrate that PCH-linked mutations of TOE1 impact many different aspects of TOE1 biochemistry, providing novel insights which may provide potential therapeutic strategies to treat PCH7 patients. In addition, these mutations provide a library of TOE1 variants that will be useful for future studies of TOE1 function and regulation.

## INTRODUCTION

Deadenylases play critical cellular roles in regulating the maturation and degradation of mRNAs and many non-coding RNAs (ncRNAs) through the cleavage of RNA poly(A) tails. Processing of RNAs by deadenylases is important for the stabilization, degradation, regulation, and/or function of the RNA substrate (1–5). Currently, there are three known human ncRNA deadenylases: poly(A) specific ribonuclease (PARN), U6 snRNA phosphodiesterase 1 (USB1), and Target of Egr1 (TOE1) (6). TOE1 is a unique and essential enzyme capable of both deadenylase (removal of Poly(A)’s) and 3’-to-5’ exonuclease activities (7). The cellular role of TOE1 is to remove the 3’-post-transcriptionally added poly(A) tails of small ncRNAs during their maturation (6, 8–12), and to protect these RNA substrates from early degradation and processing (6, 8, 12). Although many ncRNA substrates are shared among TOE1 and PARN, TOE1 is the deadenylase responsible for targeting Pol-II transcribed small nuclear RNA (snRNA) precursors (6, 8–12). Evidence of a defect in snRNA processing due to loss of TOE1 was shown by a rescue experiment utilizing a catalytically inactive mutant and wildtype TOE1 (8).

Additionally, missense mutations found throughout the *TOE1* gene have been associated with a neurological disorder called pontocerebellar hypoplasia type 7 (PCH7). Pontocerebellar hypoplasia (PCH) is an autosomal recessive pediatric neurodevelopmental and neurodegenerative disease classified based on genotype and phenotype (13, 14). Biallelic (homozygous) and compound heterozygous mutations of TOE1 are linked to (PCH7) (8, 13–18). Hallmarks of PCH7 are similar to other PCH subtypes: atrophy of the pons and cerebellum, breathing abnormalities, and muscular hypotonia; however, PCH7 also includes a unique phenotype of hypogonadism (8, 13). Although, fibroblasts derived from patients with PCH-linked TOE1 (E220K, F148Y, and A103T) mutations revealed reduced TOE1 protein levels (8), the mechanism by which TOE1 mutations lead to PCH7 remains poorly understood. Similarly, little is known about how TOE1 recognizes its preferred RNA substrates, or how it is regulated.

PCH7-linked mutations occur throughout the TOE1 protein and have been linked to decreased protein levels and changes in cellular localization (8, 18), which suggests that mutations could impact different aspects of TOE1 function and regulation. Presently, no robust biochemical analysis of these mutations has been performed. Here, we utilize AlphaFold predictions and biochemical assays to better understand how PCH7-linked mutations impact TOE1 function *in vitro* and *in vivo*, to better understand how they might impact TOE1 cellular functions. We further extrapolate what we learned from this set of mutants to provide novel insights into TOE1’s biochemical properties.

## MATERIAL AND METHODS

### Cloning and generation of plasmids

A codon-optimized gene for TOE1 and all variants was generated by Genscript. TOE1 genes were subcloned into the pHis2-parallel vector (with an N-terminal His tag) using BamHI and XhoI sites and into pCDNA3.1 vectors (with N-terminal HA, GFP, and Flag tags) using KpnI, BamHI, and HindIII sites, using standard protocols. Cloning into pCDNA vectors was performed both in house and by Genscript. A table of plasmids used in this work can be found in Table S1.

### Recombinant expression and purification of TOE1 from *E.Coli*

His-TOE1 was expressed in BL21(DE3) Rosetta II pLac(I) cells (Millipore). Cells were grown at 37°C to an optical density of 0.4-0.6, in LB, and protein production was induced by the addition of 0.1 mM IPTG at 18°C, overnight. Cells were harvested and pellets were stored at −80°C until purification. Cells were lysed via sonication in 50 mM HEPES pH 8.0, 750 mM NaCl, 10% glycerol, 5 mM MgCl_2_, 10 mM imidazole, and 5 mM BME, with the addition of a Complete EDTA-free protease inhibitor tablet (Roche). Purification was conducted using HIS-60 resin (Takara). Resin was washed first with lysis buffer and then washed with 50 mM HEPES pH 8.0, 250 mM NaCl, 10% glycerol, 5 mM MgCl_2_, 10 mM imidazole, 5 mM BME. The sample was eluted in the same buffer, with the addition of 250 mM Imidazole. Protein eluate was then loaded onto a HiTrap^TM^ Heparin HP (Cytiva), equilibrated with 50 mM HEPES pH 8.0, 250 mM NaCl, 10% glycerol, 5 mM MgCl_2_ and 1 mM DTT and eluted at approximately 750 mM NaCl, over a salt gradient. The final protein concentration was measured using a NanoDrop^C^ (Thermo Scientific). The TOE1 protein concentration was calculated using the Beer-Lamber equation, using an extinction coefficient of 53,290 M^−1^cm^−1^, which was estimated using Benchling’s protein characterization tools (19). TOE1 was never concentrated for downstream experiments.

### Differential Scanning Fluorimetry (DSF)

DSF assays were performed using protein eluted from the heparin column within 24 hours of purification. Assays were performed with 1.95 μM TOE1 in 750 mM NaCl, 50 mM HEPES, 10% Glycerol, 5 MgCl_2_, and 2X SYPRO Orange (Invitrogen). Each experimental measurement was collected and averaged from technical replicates. Each reported melt temperature was recorded from an independent protein purification and experimental replicates were performed using independent protein purifications, purified on different days. TOE1 thermal stability was measured from 4 to 95°C, with 1°C/minute ramp rate on a CFX96 (Bio-Rad) using the CFX Manager 3.1. First derivative curves were obtained, confirmed to be one state, and fit to a Gaussian in GraphPad Prism version 10 to determine the melt temperature. Statistical analysis was performed using a One-Way Anova in GraphPad Prism version 10.

### TOE1 ribonuclease assays

Ribonuclease assays were performed at 37°C in 26 mM HEPES pH 8.0, 150 mM NaCl, 2.6 mM MgCl_2_, 2% glycerol, 0.4 mM DTT, 0.8 mM Spermidine, 0.08 mg/ml bovine serum albumin, 0.8% IPGAL CA-630 (Sigma Aldrich), and murine RNase inhibitor (NEB, diluted 10,000X) with 500 nM 5’-flourescein labelled RNA substrate and 250 nM TOE1. Synthetic, 5’ fluorescently labelled RNAs were synthesized and HPLC purified from DharmaCon or Genscript. The sequence and source information for all synthetic RNAs used in this study are available in Table S2. Each set of assays included TOE1^WT^ as a positive control and TOE1^D64A,E66A^ as a negative control for ribonuclease contamination. Timepoints were quenched with loading dye containing urea. Samples were separated by electrophoresis on 15% TBE-Urea gels (Invitrogen) for 60 min at 180 V and then imaged using a ChemiDocMP (Bio-Rad). Analysis of TOE1 cleavage was performed using ImageJ. For the analysis, the RNA amounts across different regions of the gel were quantified, using the RNA ladder to determine product states. Total RNA across all regions was calculated with the bottom of the 20-nucleotide band used as the cut-off for what had not been fully deadenylated and the top of the 2-nucleotide band used to calculate the RNA that had completed cleavage.

### Structural predictions

Structural predictions were performed using AlphaFold 3 (20), using WT TOE1 as the input sequence. An overlay of the five best TOE1 dimer models is shown in Figure S1.

### Immunoprecipitation of TOE1^F148Y^ and TOE1^WT^ from HEK293T cells

HEK293T cells were obtained from ATCC (CRL-3216) and cultured in high glucose Dulbecco’s Modified Eagle Medium (DMEM, Gibco) and supplemented with 1X L-glutamine, 50 units/mL penicillin/streptomycin solution (Gibco), and 10% fetal bovine serum (FBS, Cytiva). Cells were routinely tested for mycoplasma. HEK293T cells were transfected with plasmid DNA, using FuGENE HD (Promega). Cells were harvested and washed with Phosphate-Buffered Saline (PBS) (GenClone). Cells were lysed with a buffer composed of 50 mM HEPES, pH 8.0, 5 mM MgCl_2_, 200 mM NaCl, 10% glycerol, 0.05% IGEPAL CA-630 (Sigma Aldrich), and 2 mM Phenylmethylsulfonyl fluoride (PMSF). Clarified cell lysates were incubated with Anti-HA agarose resin (Genscript) and then washed with the lysis buffer before the resin was boiled in loading dye. Western blots were performed using standard protocols. The antibodies used were: anti-MYC tag antibody (Clone 4A6, Millipore) and anti-HA (C29F4, Cell Signalling Technology). Samples were analyzed by quantifying the adjusted volume of the bands, using Image Lab (Bio-Rad). Then, the ratio of adjusted volumes for bait/prey were calculated and compared as a fraction of the wildtype bait/wildtype prey. Statistical significance was calculated using a One-Way Anova in GraphPad Prism version 10.

### Size exclusion chromatography

TOE1 samples were separated on a Superdex 200 Increase (Cytiva) equilibrated with 50 mM HEPES, pH 8.0, 250 mM NaCl, 2% glycerol, 5 mM MgCl_2_, and 1 mM DTT. These experiments were performed utilizing recombinant TOE1 purified from *E.Coli.* Assays of peak fractions were performed as described above.

## RESULTS

### Architecture of TOE1 and PCH variants utilized in our studies

To study the impact of PCH mutations on the biochemical properties of TOE1, we first selected eleven established PCH-linked missense mutations in TOE1 (8, 16), located throughout the protein (Figure 1A). We focused primarily on the initially reported missense mutations of TOE1 and a subset of others reported in another study (16). We initially included PCH mutations TOE1^A54V^ (16) and TOE1^Y231Δ^ (8), but neither purified sufficiently for further biochemical studies. As a monomer, TOE1 is a 56.5 kDa protein with three main domains: the DEDD-type deadenylase domain, a C3H zinc finger, and a nuclear localization signal (NLS) (11, 21, 22).

**Figure 1:**
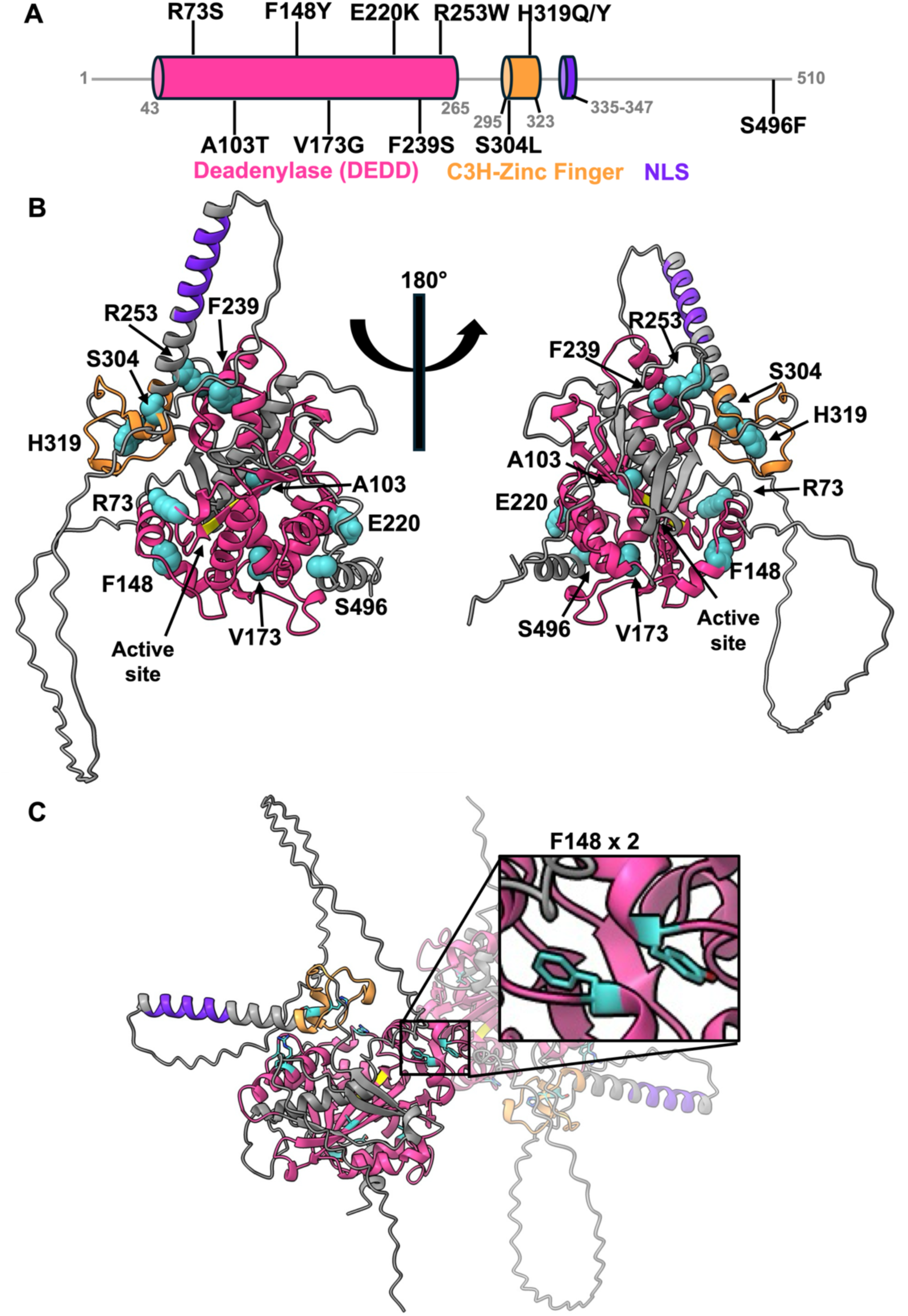
Architecture of TOE1 and location of PCH-variants. A) The domain architecture of TOE1 labeled with the location of the PCH-linked mutations covered in this work. The deadenylase domain is colored in pink, the C3H zinc finger in orange, and the nuclear localization signal (NLS) in purple. PCH-linked variants are labeled in their respective location. B) An AlphaFold generated ribbon structure of the TOE1 monomer with PCH-linked TOE1 variants colored in a teal/cyan, with two active site (D64,E66) residues highlighted in yellow. C) An AlphaFold generated ribbon structure of homodimer of TOE1, with a zoom-in for the dimer interface with F148.

We then utilized AlphaFold3 (20) to predict the structure of TOE1 and reveal the potential molecular environments in which these PCH variants occur (Figure 1B). The model reveals a globular deadenylase domain, which contains the catalytic core. The deadenylase domain is connected to the C3H zinc finger, which forms the base of a helix containing the NLS. In addition to these central ordered domains, there is a loop and the C-terminal tail, which are both unstructured. Most of the PCH-linked variants are found within the deadenylase domain with some scattered near or within the zinc finger domain (Figure 1A, B). Notably, while S496 is not within the previously defined boundaries of the deadenylase domain of TOE1 (Figure 1A), in the AlphaFold structure (Figure 1B), there are two alpha-helices and three beta-strands (which appear to extend an existing beta-sheet) that contribute to the structure of the deadenylase domain (shown in grey) and S496 resides in one of these helices. No mutations were located directly within the active site, the NLS, nor in the major disordered regions.

In addition, because cellular deadenylases frequently function as homo and heterodimers (6, 7, 23), we also used AlphaFold 3 (20) to predict the homodimerization of TOE1. The predicted TOE1 homodimer structure revealed that F148 appears at the dimer interface (Figure 1C), suggesting that this mutation could disrupt homodimerization.

### Purification of active recombinant TOE1 variants to high purity

To begin biochemically studying TOE1, we expressed TOE1 variants in *E.coli* and performed affinity purification using an encoded N-terminal His-tag, similar to previous studies (7). We then performed an additional purification using heparin affinity chromatography, which generated a concentrated fraction of TOE1 for further study (Figure S2A). To confirm that the purity of this workflow resulted in protein free from substantial ribonuclease contamination, we tested the activity of TOE1^WT^ and a previously reported catalytic-deficient variant (TOE1^D64A, E66A^) (7). We utilized an RNA substrate that allowed for visualization of both the deadenylase and exonuclease activities of TOE1^WT^ (Figure 2A). Deadenylation was observed as the cleaved RNA migration from the 40-nucleotide RNA ladder band to the 20-nucleotide ladder band. Cleavage occurring after deadenylation was classified as exonuclease activity. The RNA substrate selected for these studies was a previously published deadenylase substrate with a stem-loop with alternating weak (A:U) and strong (C:G) base pairs, an AAAU, and then a 20-nucleotide 3’ poly(A) tail (24). Use of this dual-function substrate allowed us to observe TOE1^WT^ deadenylation of the poly(A) tail and exonuclease decay of the remaining substrate, which are absent in the catalytic-deficient TOE1^D64A, E66A^ sample (Figure 2B). We also performed a control experiment in which we purchased the same RNA with the label swapped to the 3’-end and saw that TOE1 was unable to cleave the RNA, supporting the purity of our workflow and that TOE1 does not have 5’-exonuclease or endonuclease functions on the substrate tested (Figure S2B).

**Figure 2:**
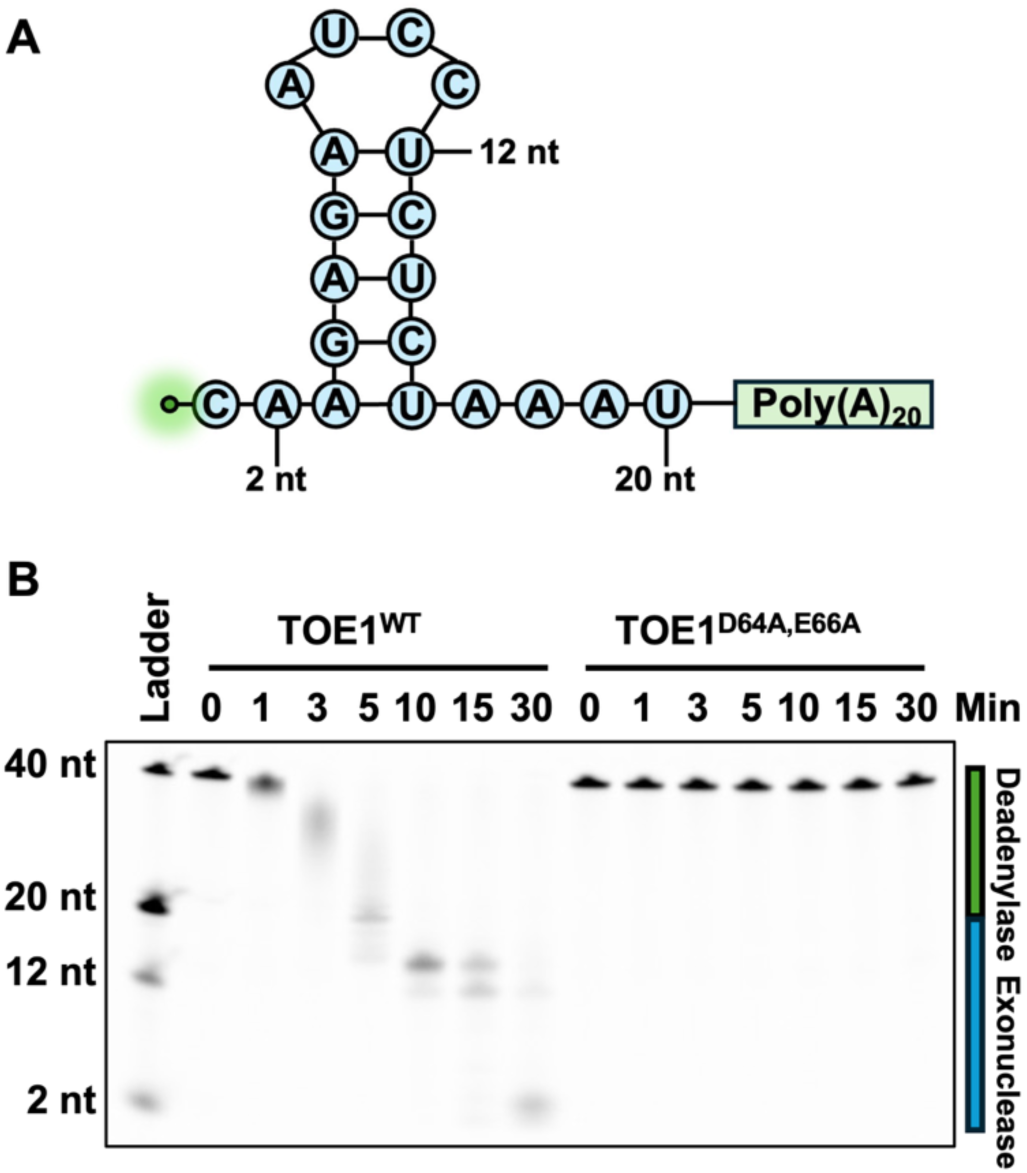
Recombinant TOE1^WT^ functions as a deadenylase and exonuclease. A) The 5’-fluorescently labeled RNA substrate used in the ribonuclease assays contains a stem loop (CAAGAGAAUCCUCUCU), follow by AAAU and a 20-nucleotide poly(A) tail (24). The regions where the RNA substrate sequence was truncated from the 3’ end for the RNA ladder controls, is marked on this image. B) A representative 30-minute time course ribonuclease assay for TOE1^WT^ and TOE1^D64A,E66A^. A green bar to the right of the gel notates the region of the gel that is deadenylase activity, while a blue bar indicates the region of exonuclease activity.

### PCH-linked variants can impact TOE1 deadenylase activity

We then utilized our ribonuclease assay to assess how PCH-linked variants could alter TOE1 activity *in vitro*. Each PCH-linked TOE1 variant was assayed for deadenylase and exonuclease activity across a time course (Figure 3). Under our assay conditions, we reproducibly saw that TOE1^WT^ would deadenylate most of the 20-nucleotide poly(A) tail of its RNA substrate in 5 minutes, with almost complete deadenylation in 10 minutes (Figure 2B, Figure 3A). We thus used the 10-minute timepoint to compare deadenylation completion across PCH variants. For this measurement, we calculated the amount of RNA that had been completely deadenylated at 10 minutes as measured against the RNA ladder. From our data, we observed statistically significant reductions in the deadenylase activity for TOE1^R73S^, TOE1^A103T^, and TOE1^F148Y^ (Figure 3B, C, D, M, and Table S3). R73, A103, and F148 along with V173, are located near the active site (Figure 4A). We also observed a reduction in the deadenylase activity of TOE1^F293S^, although it was not statistically significant. Although not highlighted by the analysis, in our gels we observed a slight increase in the deadenylase activity of the zinc finger mutations TOE1^H319Q^ and TOE1^H319Y^.

**Figure 3:**
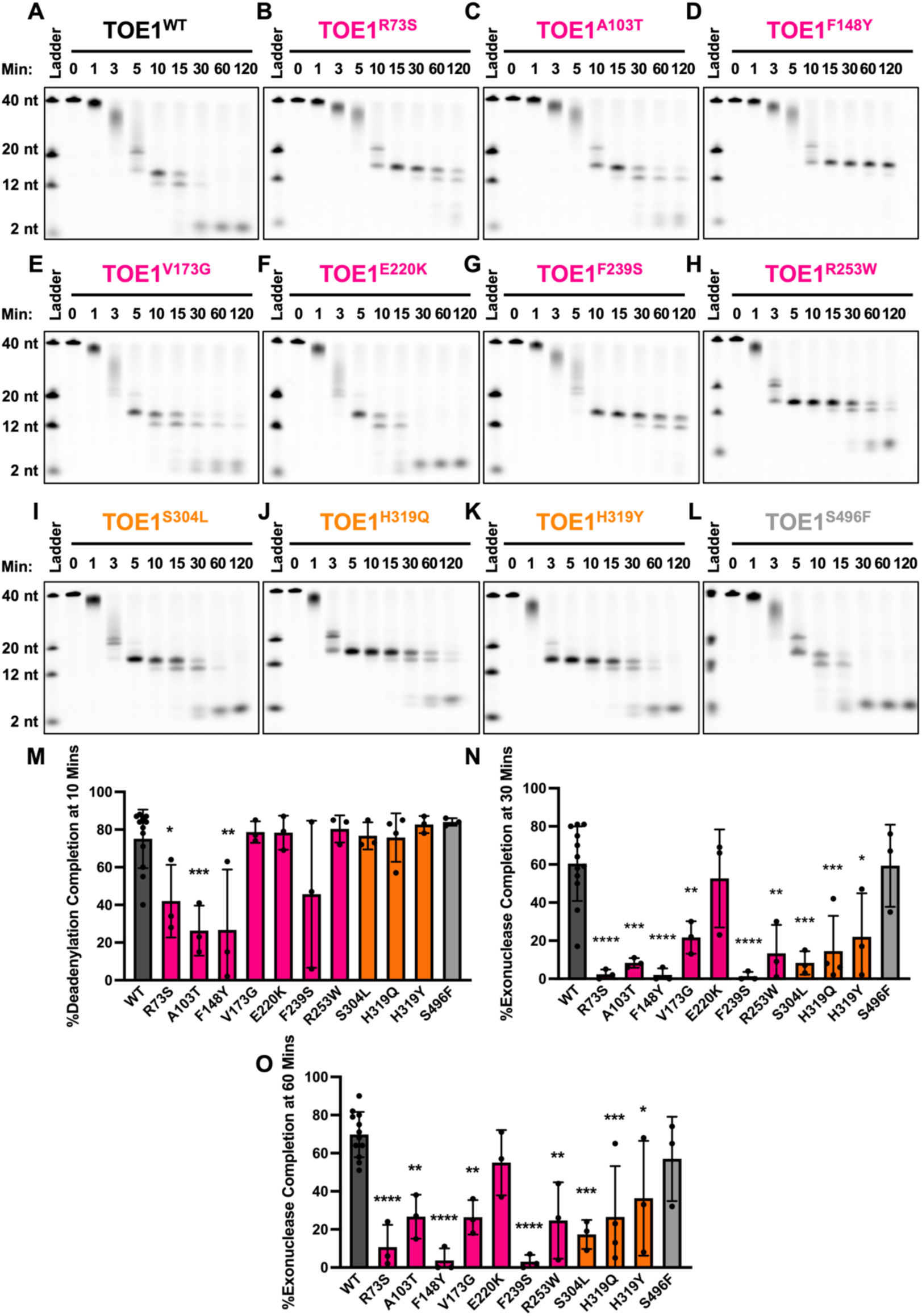
Ribonuclease assays reveal altered deadenylase and exonuclease functions for PCH-linked TOE1 variants. TOE1 (250 nM) was assayed for cleavage of a 5’-fluorescently labeled RNA substrate (500 nM). A) WT, B) R73S, C) A103T, D) F148Y, E) V173G, F) E220K, G) F239S, H) R253W, I) S304L, J) H319Q, K) H319Y, and L) S496F. M) Percent of deadenylation completed at 10 mins and N) Percent of exonuclease activity completed at 30 mins. O) Percent exonuclease activity completed at 60 mins. Each datapoint is an experimental replicate from independent protein purifications. *= p<0.05, **=p<0.01, ***=<0.001, ****=<0.0001 using a One-Way Anova with Dunnett’s multiple comparisons test.

**Figure 4:**
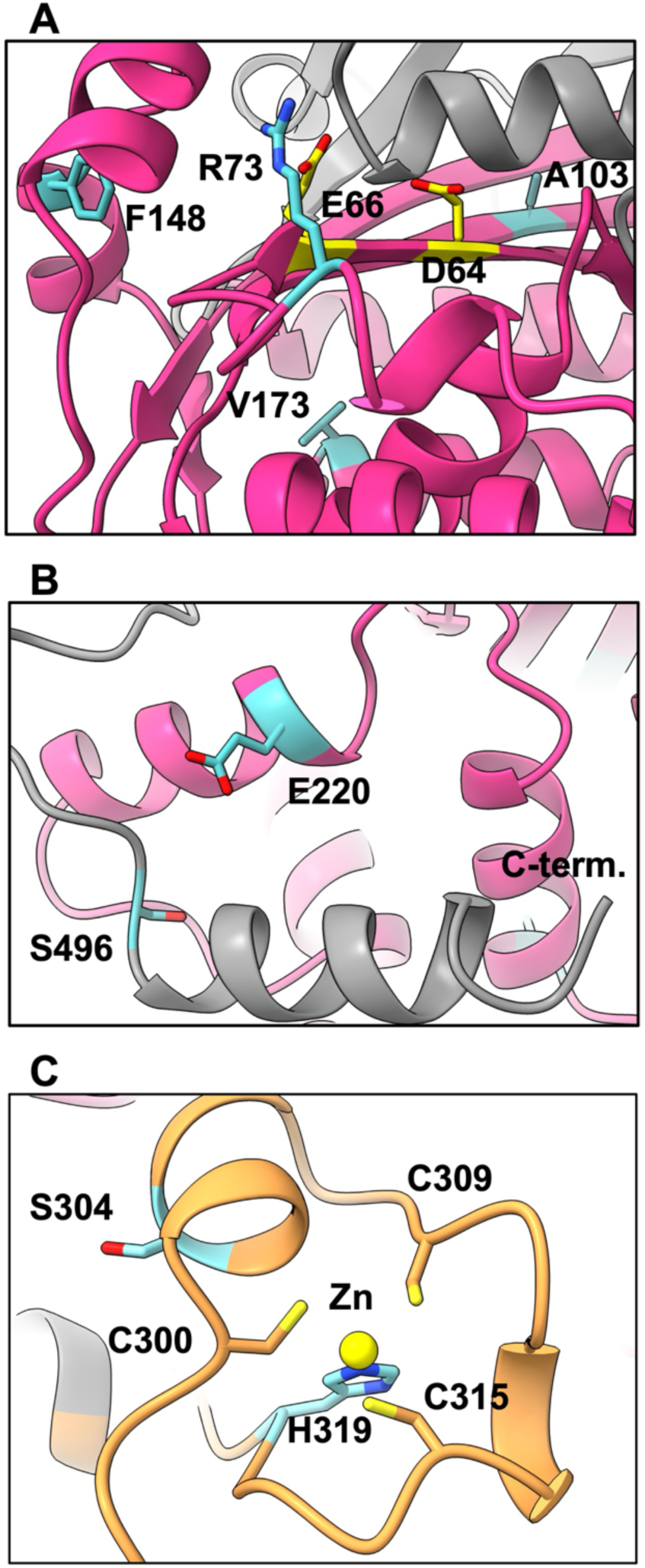
Structural predictions of TOE1 PCH mutations. Structural depiction of the molecular environment of amino acids with PCH-linked variants that had notable ribonuclease activity. Amino acids with PCH-linked mutations, highlighted in this work, appear in teal/cyan. A) Most notable impacts were seen for R73, A103, and F148 for their impact on TOE1 deadenylase activity. The catalytic residues (D64, E66) are colored yellow. B) E220 and S496, which are located near the C-terminus of TOE1, were identified for having activity similar to wildtype. C) An AlphaFold3 (20) generated TOE1 with a zinc ion (yellow) in the zinc finger domain (orange), coordinated by H319 and C300, C309, C315.

### PCH-linked variants frequently impact TOE1 exonuclease activity

Using these assays, we also examined TOE1’s exonuclease activity by observing how TOE1 cleaved the RNA substrate following deadenylation. Because wildtype TOE1 typically finished cleaving the RNA by the 30-minute timepoint (Figure 2B, Figure 3A), we selected the 30-minute and 60-minute timepoints to compare variants (Figure 3N, O). From these data, we observed that, in contrast to the deadenylase activity, only two mutants had exonuclease profiles similar to TOE1^WT^ : TOE1^E220K^ and TOE1^S496F^ (Figure 3F, L, N, and Table S3). When we examined the location of E220 and S496, we observed that they are in the same region of TOE1, connecting a C-terminal alpha helix to the deadenylase domain of TOE1 (Figure 4B), suggesting this region is not important for catalytic function but has another important role in TOE1 function or regulation.

All other mutants had statistically significant reductions in exonuclease activity, compared with TOE1^WT^, over the time course, under these assay conditions (Figure 3N, O, and Table S3), even if some of the variants had potentially faster deadenylation rates. The ability of our substrate to reveal the exonuclease defects even if there were deadenylase impacts highlights the benefit of using this substrate for these studies. Interestingly, we observed an intermediate impact on exonuclease activity by the H319Y variant. TOE1^H319Q^ and TOE1^H319Y^ generally approached completion over the time course of our assays, with TOE1^H319Q^ having a more significant impact than TOE1^H319Y^ (Figure 3J, K, N, O, and Table S3), with both TOE1^H319Q^ and TOE1^H319Y^ being slower than TOE1^WT^, TOE1^E220K^, or TOE1^S496F^ (Figure 3N, O). H319 lies within the zinc finger domain.

When we examined the sequence and structure of the zinc finger, we saw that there are two histidine residues within the C3H zinc finger domain, H295 and H319. Therefore, we used AlphaFold to model zinc into our full-length TOE1 structure. The AlphaFold predicted structure suggests that H319 is best positioned to form the zinc binding pocket with the three cysteines. This is supported by an unpublished zinc-bound NMR structure of the zinc finger of TOE1 that is deposited in the PDB (PBDID:2FC6). In both the AlphaFold (Figure 4C) and NMR structures, the zinc is coordinated in a similar manner by H319, C300, C309, and C315. Thus, we believe that H319 is the functional C3H zinc finger histidine. Because zinc fingers can play important roles in the binding and recognition of substrates by nucleic acid binding proteins (25), we hypothesize that the intermediate reduction in exonuclease activity that we observed for TOE1^H319Q/Y^ could be related to something about RNA recognition.

### PCH mutations can impact the thermal stability of TOE1

Other ribonucleases are also known to have genetic mutations linked to PCH, including the RNA exosome (13, 14, 26–30) and tRNA splicing endonuclease complex (31). Previous studies found that PCH mutations resulted in reduced protein stability (28, 30 31), suggesting that reduced protein stability may be characteristic of PCH mutations. We thus reasoned that the reduced ribonuclease activity that we observed in our ribonuclease assays could be due to reduced stability of TOE1. Therefore, we tested the thermal stability of TOE1 *in vitro,* using differential scanning fluorimetry (DSF). Thermal stability analysis revealed that TOE1^WT^ had a melting temperature (T_m_) of 53.54°C ± 0.70 and catalytically inactive TOE1 had a T_m_ of 57.11°C ± 0.32 (p <0.0001) (Figure 5A). The catalytically inactive TOE1 has two alanine mutations in its active site (D64A and E66A) which, despite blocking TOE1 activity, may contribute to an increase in thermal stability compared with active TOE1. Analysis of our eleven PCH-linked variants revealed that eight mutants (TOE1^A103T^, TOE1^F148Y^, TOE1^V173G^, TOE1^E220K^, TOE1^F239S^, TOE1^H319Q^, TOE1^H319Y^, and TOE1^S496F^) had statistically significantly lower thermal stabilities than WT, although all remain above physiological temperature. We observed the most severe reduction in T_m_ for TOE1^A103T^, TOE1^F148Y^, and TOE1^V173G^, which are all in regions that are near the active site and appear to be in regions important for TOE1 stability (Figure 5B, C). Notably, TOE1^A103T^ and TOE1^F148Y^ appear to impact both TOE1’s ribonuclease activities and thermal stability. It remains unclear if the reduction in ribonuclease activities of TOE1^A103T^ and TOE1^F148Y^ is because of their reduced thermal stability. We also found that three PCH-linked variants (TOE1^R73S^, TOE1^R253W^, and TOE1^S304L^) did not have a thermal stability that was significantly different from TOE1^WT^ (Figure 5A and Table S4), despite each of the three having reduced ribonuclease activities. Inspection of these three amino acids revealed that R73 is solvent exposed, near the active site (Figure 5D), while S304 and R253 appear to be important for stabilizing the helix containing the NLS (Figure 5E). Together, these results demonstrate that many, but not all TOE1-linked PCH mutations can impact the thermal stability of TOE1, and that thermal stability may not be the only cause of reduced ribonuclease activity.

**Figure 5:**
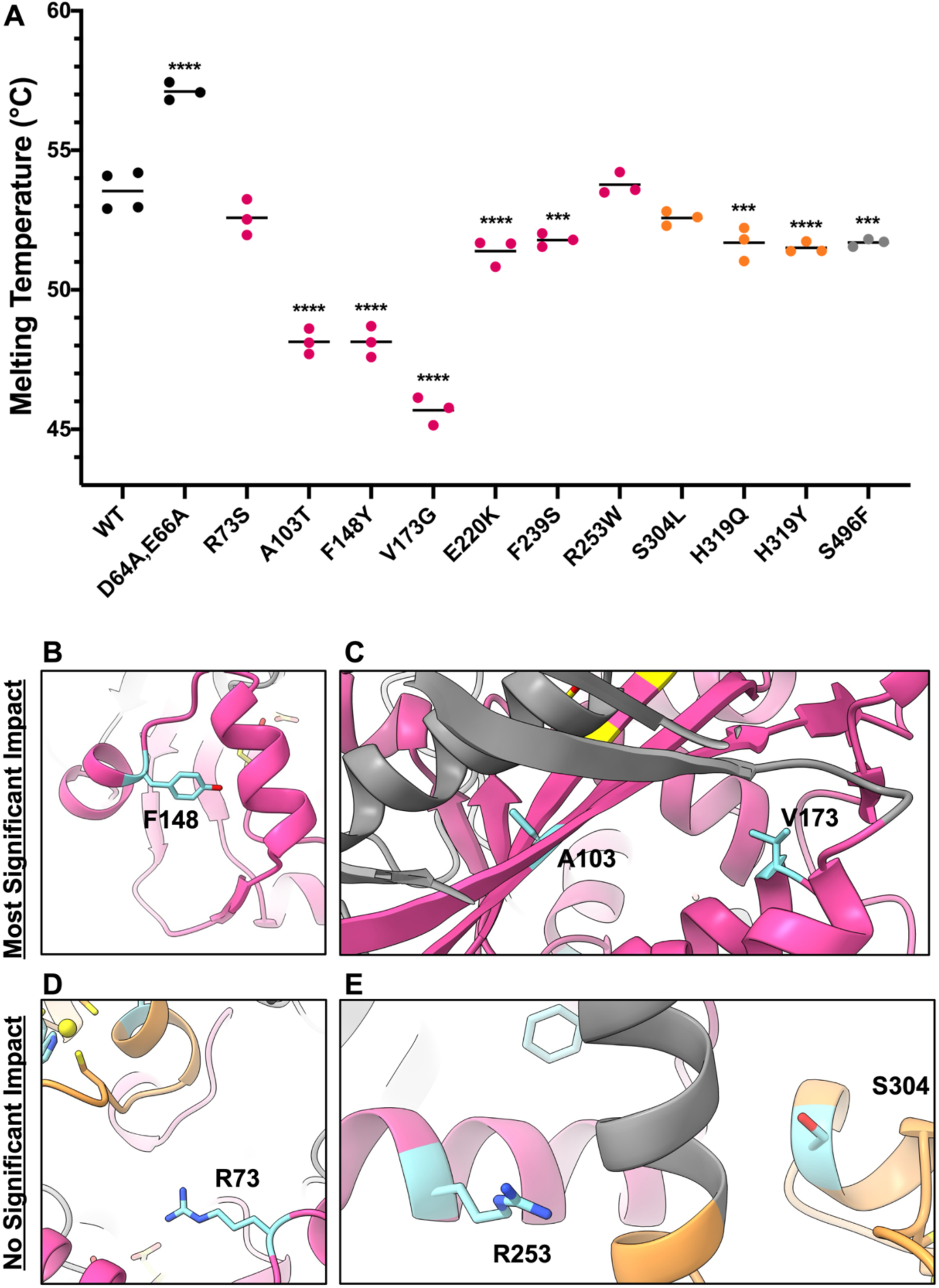
PCH mutations can impact the thermal stability of TOE1. A) Melting temperatures of TOE1 variants, calculated using DSF. Each point shown is an experimental replicate (unique protein purifications with assays on different days). Data was analyzed by One-Way ANOVA (***= p<0.001 and **** =p< 0.0001), using Dunnett’s multiple comparisons test. N=3 replicates, except for WT, for which N=4. B,C) The molecular environments of amino acids with variants that had the greatest impact on thermal stability F148, A103, and V173. D,E) The molecular environments of amino acids with variants that had no significant impact on thermal stability R73, R253, and S304. The amino acids with PCH-linked variants are shown in teal/cyan and are labeled.

### TOE1 F148Y has reduced oligomerization

Finally, we wanted to test the hypothesis that the F148Y mutation could disrupt a TOE1 homodimer. Because the oligomeric state of TOE1 has not been shown biochemically, we utilized a two-prong approach to test if TOE1 forms a homo-oligomer both *in vitro* and *in vivo*. To test if TOE1 forms a homo-oligomer *in vivo* and to assess the impact of TOE1^F148Y^, we performed TOE1 overexpression and immunoprecipitation experiments in which we co-expressed HA and MYC tagged variants of TOE1^WT^ and TOE1^F148Y^ in HEK293T cells. In these experiments, we co-transfected different combinations of “bait” HA-tagged versions of TOE1 (TOE1^WT^ or TOE1^F148Y^) and “prey” MYC-tagged TOE1 (TOE1^WT^ or TOE1^F148Y^). The samples were immunoprecipitated using anti-HA agarose resin to test for the interaction of HA-tagged TOE1 with either TOE1^WT^ or TOE1^F148Y^ MYC-tagged TOE1 variants. We observed the strongest interaction for TOE1^WT^ with itself (Figure 6A, lane 4 compared with lanes 2,3,5, Figure 6B) and observed reduced binding for TOE1^F148Y^/TOE1^WT^ combinations (Figure 6A, lane 2 and 3 compared with lane 4, Figure 6B). An immunoprecipitation of TOE1^F148Y^ with itself showed consistently reduced interactions (Figure 6A, lane 5 compared to lanes 2-4, Figure 6B). These data support the idea that TOE1 forms a higher order oligomer *in vivo* and that the TOE1^F148Y^ mutation can disrupt this oligomerization.

**Figure 6:**
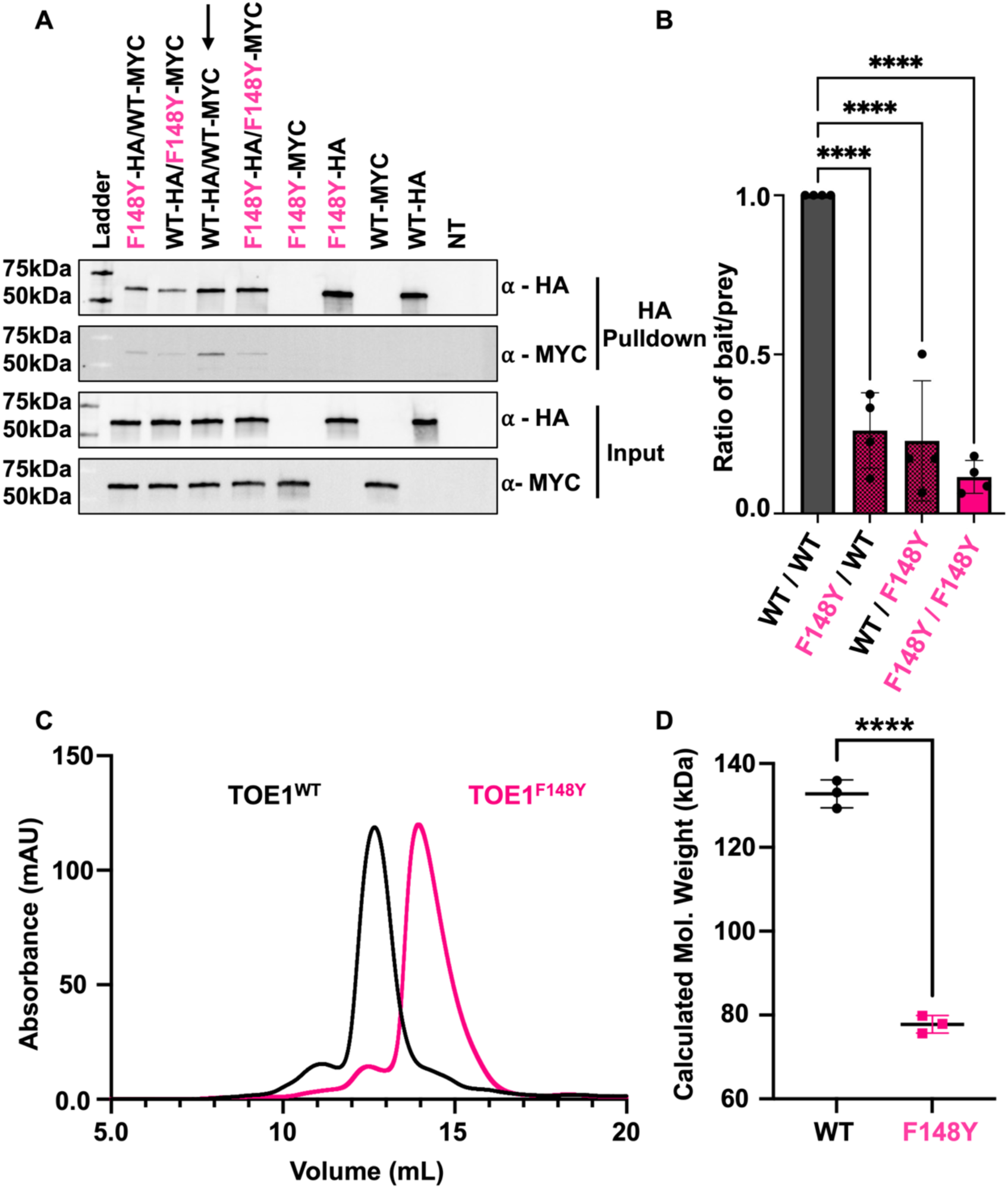
F148Y impairs TOE1 oligomerization. A) Immunoprecipitation experiments of TOE1^WT^ (black) and TOE1^F148Y^ (pink) tagged variants. An arrow notes the WT/WT sample. B) The normalized ratio of the intensity of the bait TOE1 band/ intensity of the prey TOE1 band (N= 4 independent experiments), normalized to the WT/WT ratio for each set of experiments. **** =p <0.0001 as analyzed by One-Way Anova using the Tukey’s multiple comparisons test. C) Size exclusion chromatography graph of recombinant TOE1^WT^ (black) and TOE1^F148Y^ (pink). D) Calculated molecular weight (MW) for TOE1^WT^ (black) and TOE1^F148Y^ (pink), each datapoint represents an independent protein purification. **** =p <0.001 as analyzed by an unpaired two-tailed t-test using Welch’s correction.

To test TOE1 oligomerization by another method, we utilized size-exclusion chromatography to test the state of our recombinant TOE1, from bacteria, and found that TOE1^F148Y^ elutes later than TOE1^WT^ (Figure 6C). Based on the calibration of our column, we calculated an approximate size for TOE1^WT^ at 132.8 ± 3.3 kDa (Figure 6D) which suggests that TOE1 is forming a homodimer, while the molecular weight of TOE1^F148Y^’s elution volume, 77.8 ± 2.1 kDa, aligns with a TOE1 monomer.

We also tested all the other TOE1 variants, at least once, and did not observe any other variant that impacted TOE1’s oligomeric state on the sizing column (Figure S3). We confirmed the activity for TOE1 in each peak eluate fraction of TOE1^WT^ and TOE1^F148Y^ (Figure S4A, B) and for the other TOE1 variants (N≥1). We observed an increase in activity after passing each TOE1 variant through the sizing column (Figure S4), compared to the heparin samples, in equivalent assay conditions, but the overall trends in ribonuclease activity, compared with TOE1^WT^ did not change.

Therefore, our results suggest that TOE1 forms a homodimer and that TOE1^F148Y^ causes TOE1 to monomerize. Because TOE1^F148Y^ retained activity after the sizing column, we conclude that dimerization is likely not essential for deadenylase activity *in vitro,* although it may still be important for overall function as we observed significant reductions in both deadenylase and exonuclease activities. It is unclear if the reduction we observed is due to changes in TOE1 dimerization or other impacts from the F148Y mutation.

## DISCUSSION

Our work demonstrates that PCH-linked TOE1 mutations impact a variety of different biochemical properties of TOE1, including the *in vitro* thermal stability, ribonuclease activity of TOE1, and oligomeric state (Figure 7). Importantly, we found that most PCH variants impact the exonuclease activity of TOE1 in a more significant manner, as compared to deadenylase activity, suggesting that there could be important distinctions in TOE1’s RNA processing for patients carrying these mutations. We also found that eight PCH variants impact the thermal stability of TOE1 at a statistically significant level and determined that F148Y causes a dramatic shift in TOE1’s oligomeric state – from dimer to monomer. The key biochemical findings were strengthened by combining our experimental data with AlphaFold structural predictions to help interpret how the biochemical changes were impacted within the probable molecular environments of these mutations. These various mutations, and their varied impacts on TOE1 function, provide a library from which to better understand the overall function and regulation of TOE1 and to interpret the impact of clinically identified mutations.

**Figure 7:**
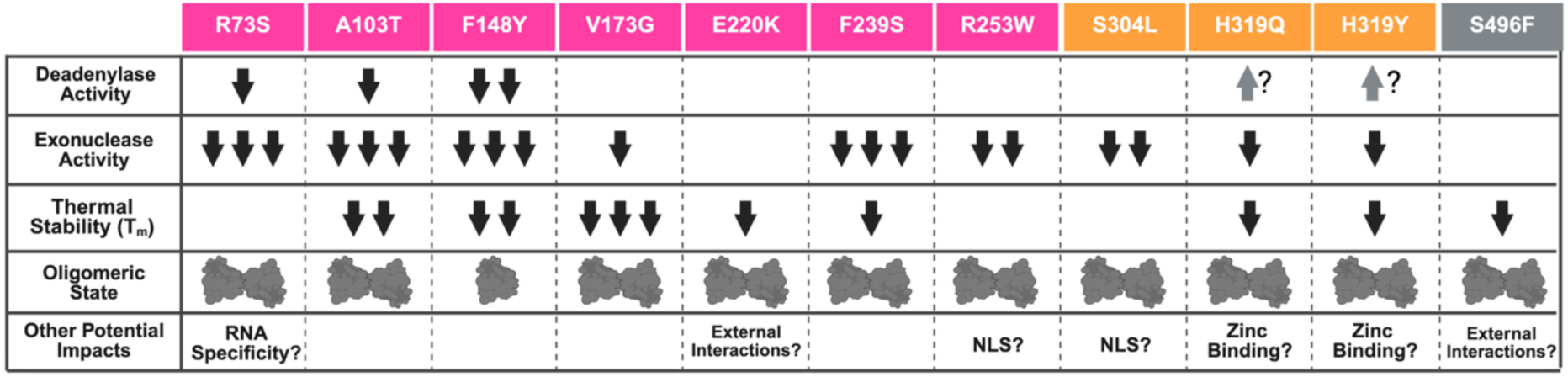
Summary of Biochemical Impacts of PCH-linked TOE1 Variants. Summary graphic of the impacts of PCH-lined TOE1 mutations on TOE1 deadenylation, exonuclease activity, stability, oligomerization, and other hypothesized impacts. Created in BioRender. Hayne, C. (2026) https://BioRender.com/7or78kz

This work builds on previous studies which have linked these and other genetic mutations in TOE1 to PCH (8, 15–17, 32). A few studies have hinted at mechanisms by which mutations may disrupt TOE1 function by examining a single characteristic of TOE1 (8, 18); those studies typically suggest differences in protein stability or protein levels. However, no additional formal experimental investigations have been published to characterize how PCH-linked point mutations impact TOE1 function, particularly *in vitro*. In addition, trying to determine how mutations impact TOE1 biology *in vivo* is difficult because only a subset of mutations have been identified as homozygous mutations (A103T, F148Y, E220K) (8), while the rest of the patients are compound heterozygotes (8, 15–18, 32). Because PCH is an autosomal recessive disease, each mutation must be causative at some level for patients to have disease, though no reports on the phenotypic severity of the different mutations have been published. Thus, studying compound heterozygous mutations in patient samples is confounded by the presence of two different mutations; however, the biochemical approaches utilized here provide a framework to decipher the underlying mechanisms by which individual mutations impact TOE1 function, stability, and oligomeric state.

Furthermore, our approach has allowed us to decipher aspects of TOE1 biology not revealed from studying patient samples alone. For example, for the biallelic mutation, E220K, patient samples revealed a reduction in TOE1 levels (8). Using our approach, we were able to conclude that E220K does not cause a major protein stability defect *in vitro*. Instead, when we examined the molecular environment in which E220 resides, we saw that E220 and S496 appear in a similar region (Figure 4B), and that based on the predicted structure, these two amino acids may form networks that hold a small predicted alpha helix in place. Although we could not clearly decern the importance of this alpha helix, we hypothesize it may be relevant to external interactions for TOE1—such as protein: protein interactions, due to being both surface exposed and away from other key regions (Figure 7). Several previous studies suggest TOE1 interaction partners (7, 8, 11, 21, 33, 34), but the molecular interface where these interactions occur is not known, which is important for validating the specificity of such interactions. As novel TOE1 binding partners are validated, this region should be assessed to see if altering these amino acids can change protein interactions.

Another aspect of TOE1 interactions that has not been elucidated is how TOE1 recognizes RNA: particularly the difference in how TOE1 recognizes and processes poly(A) and non-poly(A) substrates. Presumably, TOE1 relies on some form of poly(A) tail recognition, as has been identified for other deadenylases (4, 35, 36). Since TOE1 is also an exonuclease, how it recognizes diverse substrates is likely complex. Previous work on the similar PAN deadenylase complex, as well as other deadenylases (4, 35), has shown that zinc finger domains are important for RNA interactions (37). Interestingly, zinc fingers have been shown to impact RNA recognition and processing by deadenylases (38). In addition, a previous study of TOE1 highlighted that both the zinc finger and the NLS may play roles in RNA binding (11). Presumably the RNA binding interactions of the NLS come from the arginine rich region (RR) within the NLS as RR regions have been shown to enhance RNA binding when located next to a zinc finger (39). Since our AlphaFold model suggested that both R253 and S304 stabilize the NLS, we hypothesize that the TOE1^R253W^ and TOE1^S304L^ and the zinc finger TOE1^H319Q^ and TOE1^H319Y^ mutations specifically impact exonuclease activity due to changes in TOE1’s RNA specificity.

In the absence of an RNA-bound TOE1 structure, further studies will be needed to investigate the role of the zinc finger in TOE1 function. Our work suggests that instead of performing biochemical truncations, which might destabilize a large portion of TOE1, additional studies could be performed with a TOE1 variant that impacts the function in question. For instance, a follow-up study for RNA specificity could use the TOE1^H319Q/Y^ variants rather than removing the entire zinc finger domain. Additionally, further studies are needed to assess if the dimer has a role in RNA recognition and specificity as such a role could explain why the TOE1^F148Y^ mutation had a significant impact on both deadenylation and exonuclease activity, while still maintaining some activity. We believe that the PCH mutations tested here provide a foundation for further interrogation of TOE1 RNA substrate recognition.

Another insight gained from combining our structural prediction insights with experimental results was that some PCH-linked mutations, including TOE1^S304L^, TOE1^R253W^, and potentially TOE1^F239S^ could impact TOE1 localization (Figure 7). If these mutations were present, we hypothesize that in addition to ribonuclease changes, there could also be a decrease in nuclear localization of TOE1. TOE1 is usually found in the Cajal bodies (7, 11, 34), the subcompartment of the nucleus where most snRNA processing occurs, although there is speculation TOE1 can shuttle between the Cajal bodies and cytoplasm (12). Indeed, a recent study showed that a newly identified PCH-linked mutation, the truncated TOE1 variant TOE1^337Δ^, causes a slight localization of TOE1 to the cytoplasm when overexpressed (18), while the compound heterozygous mutation it was identified with, TOE1^F303C^, does not (18). TOE1^F303C^ is located within the zinc finger and can presumably impact ribonuclease activity. The approaches we outlined here could be used in the future to assess novel and addition mutations, such as TOE1^F303C^.

This work should be interpreted with the obvious limitation that while our *in vitro* studies are able to provide unique insights, they may not fully recapitulate the environment or protein levels of TOE1 in cells, nor the physiological substrates of TOE1, which will carry various modifications. In addition, it is unclear how post-translational modifications could support or alter TOE1 stability *in vivo* or what differences in thermal stability should be considered physiologically relevant. Nevertheless, we believe it would be useful to apply the experimental approaches we optimized here to additional TOE1 variants, to better understand if cellular changes are the result of biochemical properties or some additional feature. Furthermore, we believe the mutations assayed here will be key variants for further investigation of TOE1 function and regulation.

The work presented here contributes to the field by providing the first detailed characterization of PCH mutations, and a biochemical assessment of their impact on TOE1’s distinct ribonuclease activities. We present the first *in vitro* thermal stability data for TOE1 and have determined conditions compatible with size exclusion chromatography, which were not previously reported, allowing us to determine that TOE1 is likely a dimer. These biochemical characterizations provide key insights and the groundwork for future studies while also providing the foundation for the development of customized therapeutics targeting specific aspects of TOE1 biology. For example, our findings suggest previously measured reductions in TOE1^E220K^ protein levels may not be due to a thermal stability difference, but rather due to another property, and that TOE1^F148Y^ is a monomer. The results of this work also provide a library of variants that can now be used as controls or point mutations to study different aspects of TOE1 biology, such as differentiating between TOE1 deadenylation and exonuclease activities, understanding the role of the zinc finger, oligomerization, and probing the role of the E220/S496 region in TOE1: protein interactions.

## DATA AVAILABILITY

The raw data underlying this article will be made available upon request to the corresponding author.

## SUPPORTING INFORMATION

This article contains supporting information (figures and tables) as well as additional supporting information.

## ACKNOWLEDGEMENTS

We acknowledge Phoebe Rice, Joe Piccirilli, and Viktoriya Kalinina for thoughtful feedback of the manuscript.

## SUPPORTING INFORMATION

**Table S1:** Plasmids used in this work.

| Plasmid Backbone | Insert | Source |
| --- | --- | --- |
| pHis2 | WT<br>D64A, E66A<br>R73S<br>A103T<br>F148Y<br>V173G<br>E220K<br>F239S<br>R253W<br>S304L<br>H319Q<br>H319Y<br>S496F | Original cDNA synthesis/mutagenesis:<br>Genscript<br>Subcloning: In-house |
| pCDNA3.1 | HA - TOE1 | Genscript |
|  | HA - TOE1 (F148Y) | Subcloned in house. |
|  | MYC - TOE1 |  |
|  | MYC - TOE1 (F148Y) |  |

**Table S2:**
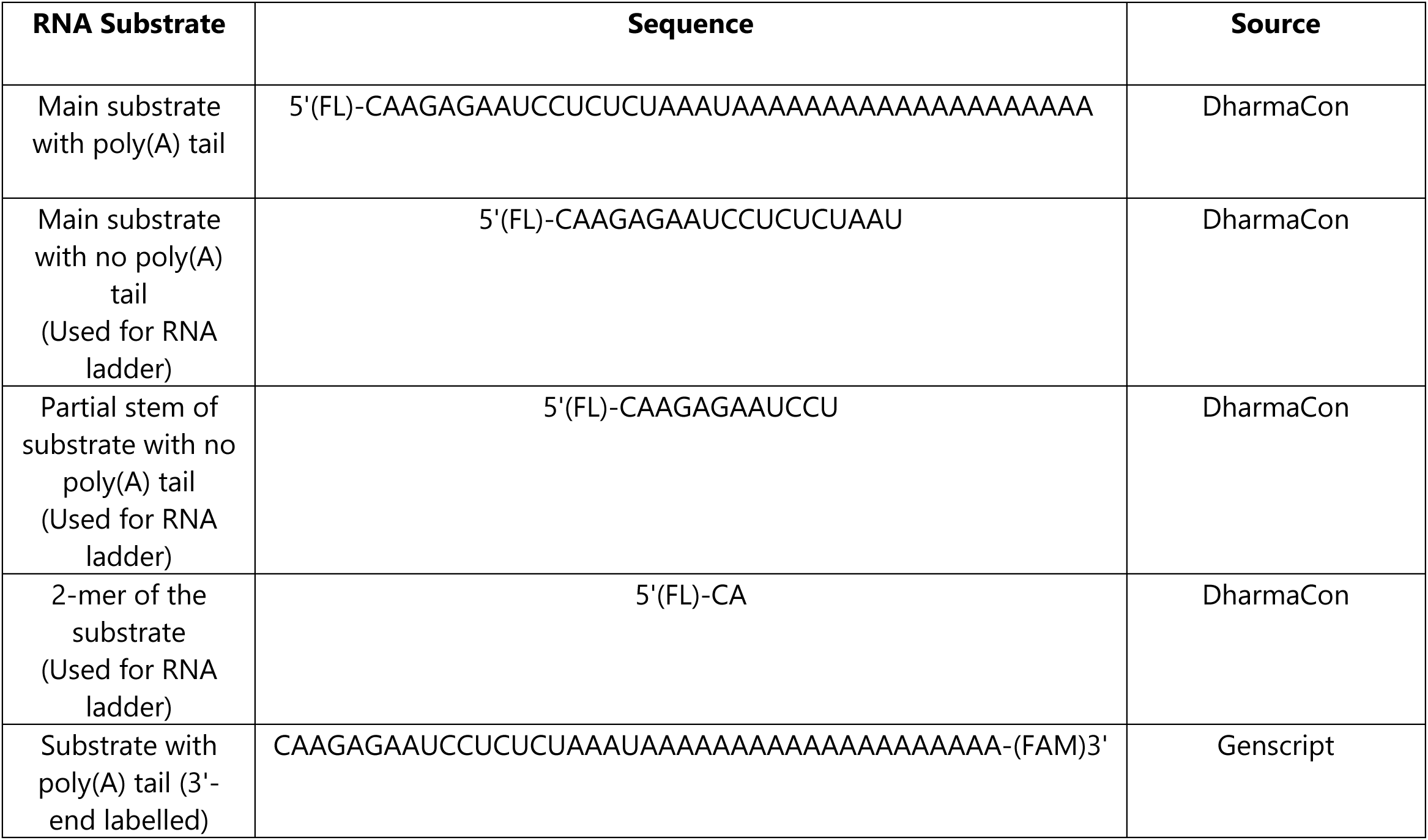
Synthetic RNA substrates used in this work.

**Table S3.** Deadenylation and exonuclease completion of TOE1 variants. Calculated percentage of substrate deadenylated at 10 minutes or percentage of substrate degraded at 30 and 60 minutes for each reported PCH mutation. Replicates (N) are from individual protein preps. Standard deviation and p-value based on a One-Way Anova are reported.

|  | N | Completed Deadenylation at 10 Minutes |  |  | Completed Exonuclease at 30 Minutes |  |  | Completed Exonuclease at 60 Minutes |  |  |
| --- | --- | --- | --- | --- | --- | --- | --- | --- | --- | --- |
|  |  | Mean (%) | Std. Dev. | p-value | Mean (%) | Std. Dev. | p-value | Mean (%) | Std. Dev. | p-value |
| **WT** | 12 | 75.08 | 15.52 | N/A | 60.42 | 19.51 | N/A | 69.75 | 11.89 | N/A |
| **R73S** | 3 | 42.00 | 19.29 | 0.0480 | 2.33 | 2.52 | <0.0001 | 10.67 | 11.72 | <0.0001 |
| **A103T** | 3 | 26.33 | 13.32 | 0.0010 | 8.33 | 2.52 | 0.0003 | 26.67 | 11.50 | 0.0023 |
| **F148Y** | 3 | 26.67 | 32.13 | 0.0011 | 2.00 | 3.46 | <0.0001 | 3.67 | 6.35 | <0.0001 |
| **V173G** | 3 | 78.67 | 5.69 | >0.9999 | 21.67 | 8.51 | 0.0094 | 26.33 | 9.07 | 0.0021 |
| **E220K** | 3 | 78.33 | 9.02 | >0.9999 | 52.67 | 25.74 | 0.9978 | 55.00 | 17.09 | 0.8067 |
| **F239S** | 3 | 45.67 | 39.02 | 0.1057 | 1.33 | 2.31 | <0.0001 | 3.00 | 3.61 | <0.0001 |
| **R253W** | 3 | 80.33 | 7.23 | >0.9999 | 13.33 | 14.98 | 0.0010 | 24.67 | 20.03 | 0.0013 |
| **S304L** | 3 | 76.67 | 7.23 | >0.9999 | 8.33 | 6.11 | 0.0003 | 17.33 | 7.64 | 0.0002 |
| **H319Q** | 4 | 75.75 | 12.84 | >0.9999 | 14.50 | 18.50 | 0.0003 | 26.50 | 26.70 | 0.0005 |
| **H319Y** | 3 | 82.67 | 4.51 | 0.9986 | 22.00 | 22.91 | 0.0102 | 36.33 | 30.14 | 0.0287 |
| **S496F** | 3 | 84.00 | 2.00 | 0.9945 | 59.33 | 21.55 | >0.9999 | 57.00 | 22.11 | 0.9070 |

**Table S4.**
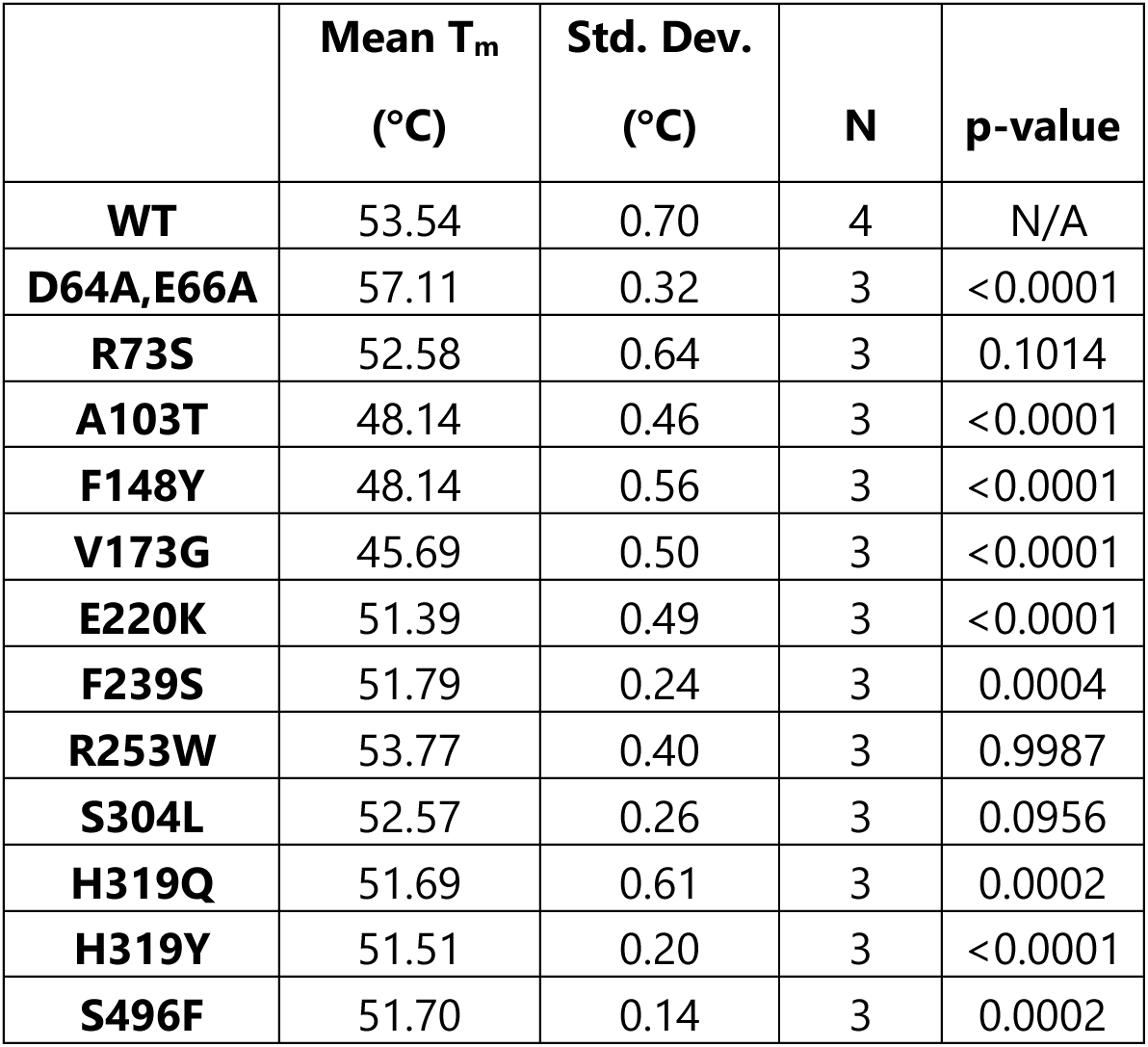
Summary table of thermal stability data. Calculated average melt temperature for each PCH mutation. Replicates (N) are from individual protein preps. Standard deviation and p-value based on a One-Way Anova are reported.

|  | **Mean T<sub>m</sub>**<br>**(°C)** | **Std. Dev.**<br>**(°C)** | **N** | **p-value** |
| --- | --- | --- | --- | --- |
| **WT** | 53.54 | 0.70 | 4 | N/A |
| **D64A,E66A** | 57.11 | 0.32 | 3 | <0.0001 |
| **R73S** | 52.58 | 0.64 | 3 | 0.1014 |
| **A103T** | 48.14 | 0.46 | 3 | <0.0001 |
| **F148Y** | 48.14 | 0.56 | 3 | <0.0001 |
| **V173G** | 45.69 | 0.50 | 3 | <0.0001 |
| **E220K** | 51.39 | 0.49 | 3 | <0.0001 |
| **F239S** | 51.79 | 0.24 | 3 | 0.0004 |
| **R253W** | 53.77 | 0.40 | 3 | 0.9987 |
| **S304L** | 52.57 | 0.26 | 3 | 0.0956 |
| **H319Q** | 51.69 | 0.61 | 3 | 0.0002 |
| **H319Y** | 51.51 | 0.20 | 3 | <0.0001 |
| **S496F** | 51.70 | 0.14 | 3 | 0.0002 |

**Figure S1:**
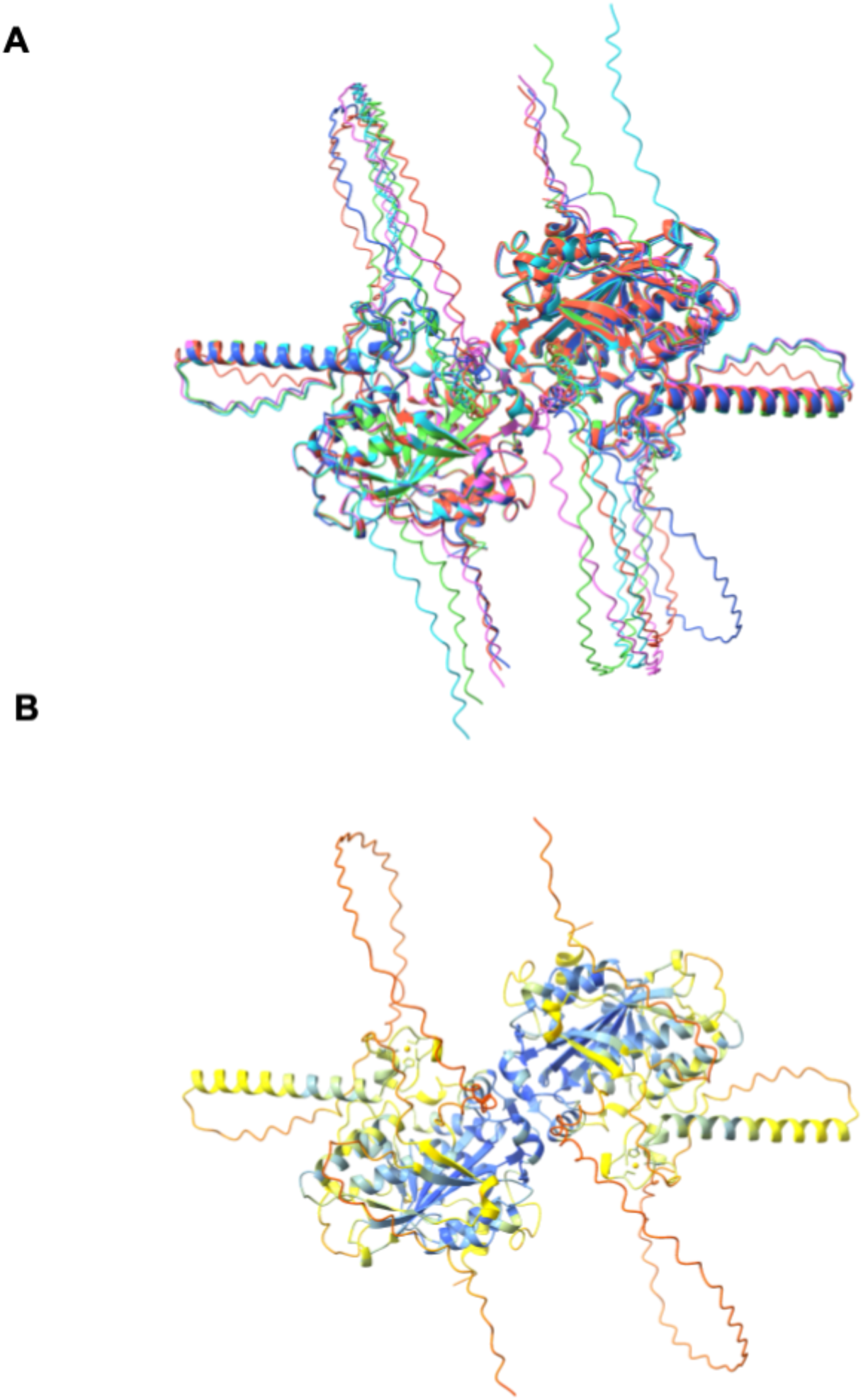
AlphaFold models of TOE1 dimer. A) The top 5 models for the TOE1 dimer modeled using AlphaFold3 are shown in an overlay. B) The best AlphaFold model is colored by pLDDT values, using AlphaFold defaults.

**Figure S2:**
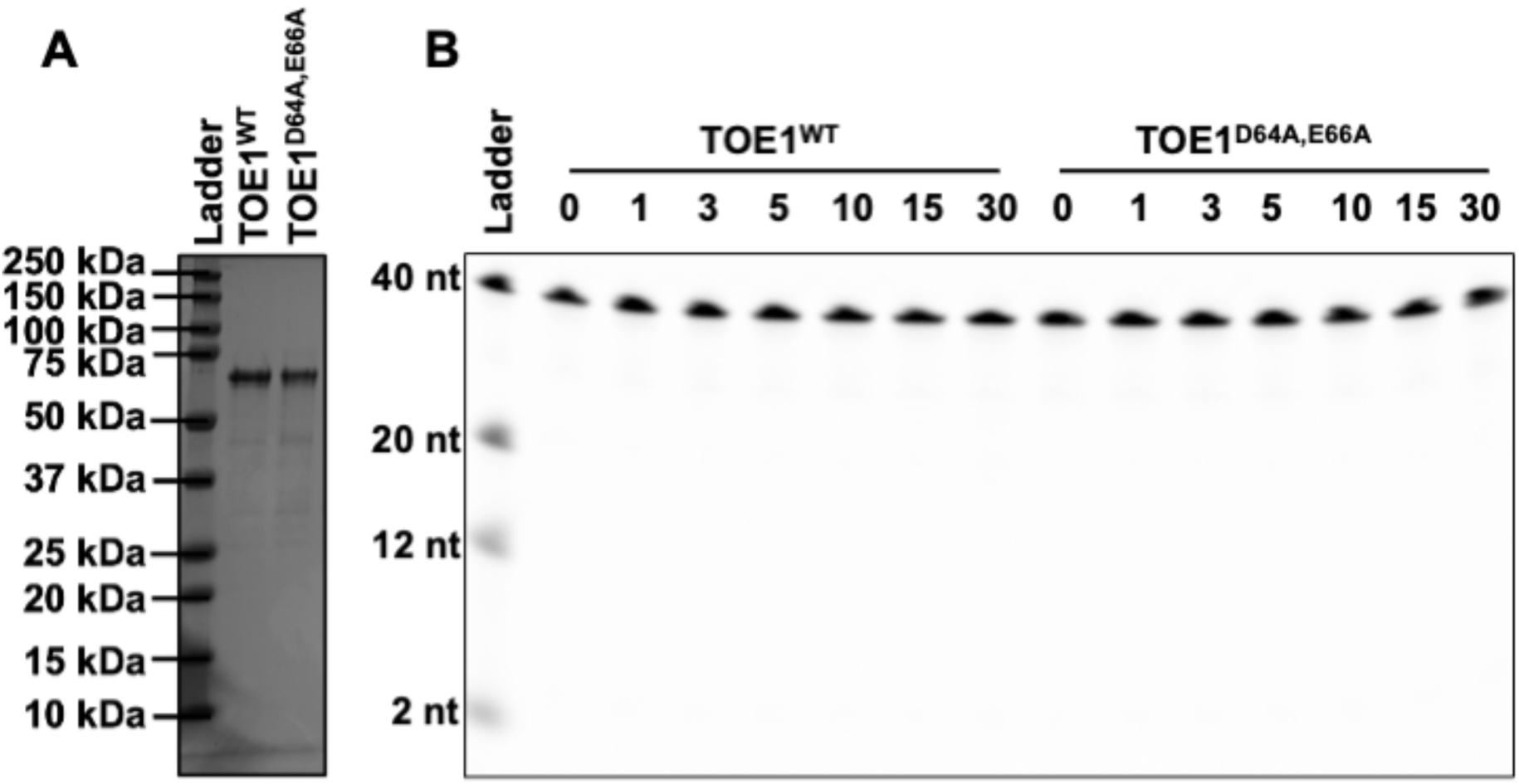
Purification of recombinant TOE1. A) Representative SDS-PAGE gel of purified TOE1^WT^ and TOE1^D64A,E66A^. B) TOE1 (250 nM) was assayed for cleavage of a 3’-FAM labeled RNA substrate (500 nM) and was unable to cleave the RNA when the 3’-end is labelled with a flurophore.

**Figure S3:**
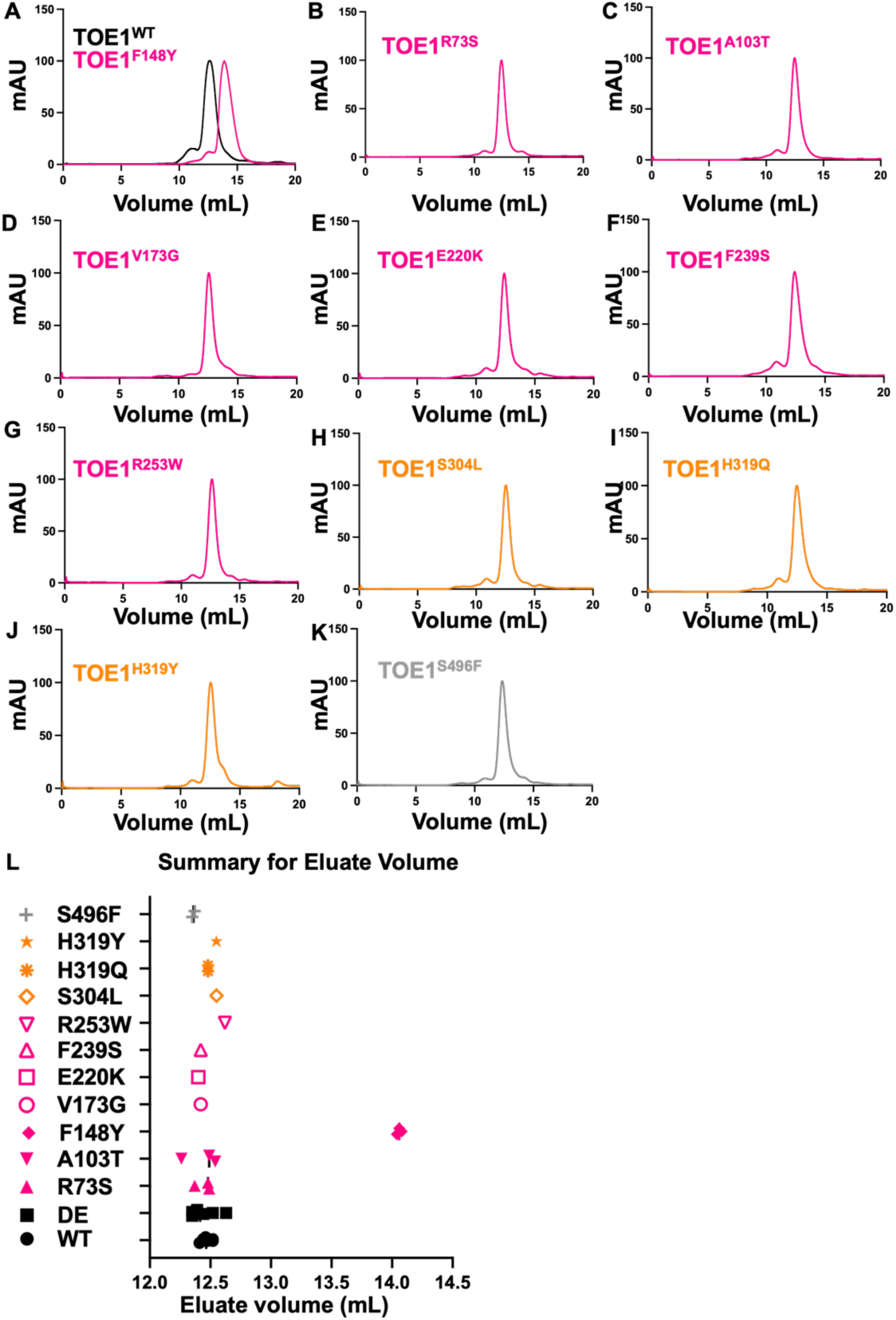
S200 gel filtration traces for PCH7-linked TOE1 variants. Profiles of size exclusion chromatographs, normalized to the highest absorbance for each run. A) F148Y and WT replicate, B) R73S C) A103T, D) V173G, E) E200K, F) F239S, G) R253W, H) S304L, I) H319Q, J) H319Y, K) S496F, L) Summary of the peak eluate volume for each PCH7-variant measurement. Each datapoint is an independent protein purification.

**Figure S4:**
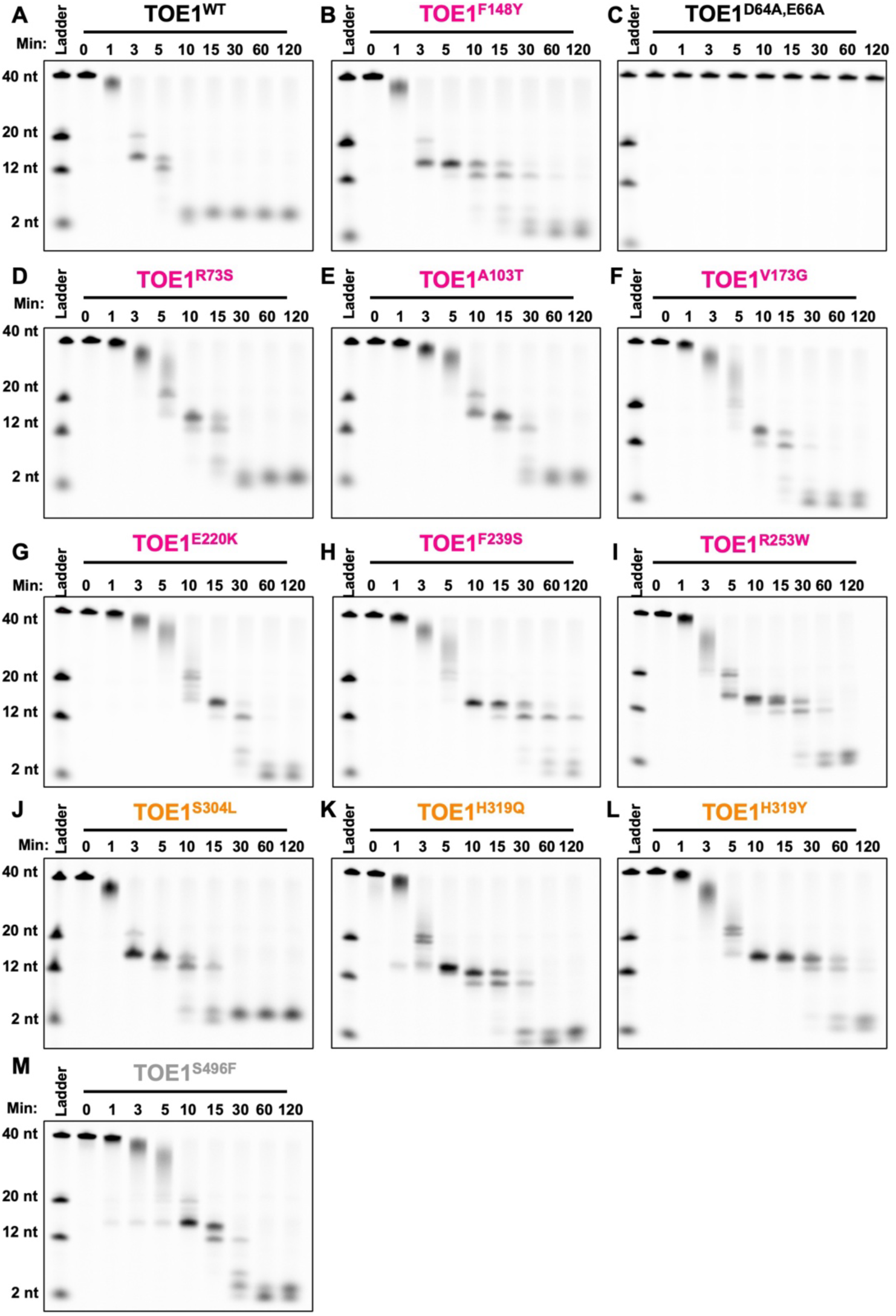
Ribonuclease assays of eluates from the gel filtration for tested variants. TOE1 (250 nM) was assayed for cleavage of 5’-fluorescently labeled RNA substrate (500 nM). A) WT, B) F148Y, C) D64A,E66A, D), E) R73S, F) A103T, G) V173G, H) E220K, I) R253W, J) S304L, K) H319Q, L) H319Y. and M) S496F.

